# Impact of the maternal environment on cardiovascular features of the offspring in a mouse model of Marfan syndrome

**DOI:** 10.1101/2025.09.10.675335

**Authors:** Isaac Rodríguez-Rovira, Cleo Geeraerts, Aleksandra-Mas-Stachurska, Ana Paula-Dantas, Bart Loeys, Victoria Campuzano, Gustavo Egea

## Abstract

**Background:** Marfan syndrome (MFS) is a systemic disorder of the connective tissue caused by heterozygous mutations in the *FBN1* gene, which encodes fibrillin 1, a glycoprotein that constitutes elastic fibers. MFS does not exhibit sexual dimorphism regarding prevalence; however, it remains unknown whether the paternal or maternal inheritance of the *FBN1* mutation affects the cardiovascular pathology of the offspring. In this study, we aimed to determine the impact of the parental origin of the *FBN1* mutation on the cardiovascular manifestations of the offspring using the *Fbn1*^*C1041G/+*^ mouse model of MFS.

**Methods and Results:** Four experimental groups were generated by crossing wild-type (WT) and MFS mice to obtain WT and MFS offspring from either a paternal (MFS-P) or maternal (MFS-M) MFS parent. Cardiovascular phenotyping of offspring was performed from childhood to adulthood (from one to six months of age), including echocardiography, tail-cuff plethysmography, histopathology, and canonical (pSmad2) and non-canonical (pERK) TGF-β signaling activity in the aortic tissue. At one month of age, both WT and MFS offspring from MFS-M presented lower body weight than those from MFS-P. However, with age, MFS-M offspring became persistently and significantly overweight. Both MFS-P and MFS-M offspring exhibited a significantly increased aortic root diameter compared with WT offspring; however, this enlargement appeared earlier in MFS-M than in MFS-P offspring. These parental and age-related differences in aortic root diameter were accompanied by increased canonical and non-canonical TGF-β signaling. The cardiac ejection fraction was reduced at early ages in both WT and MFS offspring from MFS-M compared with MFS-P, with the difference persisting only in MFS-M offspring at adulthood. Systolic blood pressure was initially lower in MFS-M offspring across both genotypes. However, it progressively increased, resulting in elevated levels in both WT and MFS offspring from MFS-M by six months of age.

**Conclusion:** Our results indicate the existence of a gestational maternal MFS environmental factor with an early impact on the aorta and heart of MFS offspring. In adulthood, this becomes normalized in the aorta but not in the heart.

## INTRODUCTION

Marfan syndrome (MFS) is caused by heterozygous genetic variants in the fibrillin-1 gene (*FBN1*) and is inherited as an autosomal dominant disorder. Twenty-five per cent of patients with MFS exhibit *de novo* mutations (1). Depending on the type of genetic variant, the disease could be potentially classified as haploinsufficiency or as dominant-negative (2,3)). Over 3,000 genetic variants have been described in *FBN1*, spanning the entire exome. However, establishing genotype-phenotype correlations has been hindered by observed inter- and intrafamilial variations in clinical manifestations, where target organs are affected to varying degrees. These variations reflect the complex and numerous functions of fibrillin-1 (4,5), and the establishment of these genotype-phenotype correlations is ongoing (3,6,7).

With respect to sex, no differences have been reported in terms of the prevalence of MFS in men and women (1,8,9), and current pharmacological treatment (mainly angiotensin II receptor blockers and β-blockers) makes no distinction between the sexes. Similar extra-aortic clinical manifestations occur both in men and women, whereas the main outcome that compromises patients’ lives—thoracic aortic aneurysm (TAA)—exhibits some differences between sexes. In the non-heritable aneurysm condition, different studies have reported a fourfold higher prevalence of subclinical aortic aneurysms in males, together with higher mortality rates in men than in women. However, women with TAA have higher mortality rates after surgical repair than men after one year of follow-up. Thus, in general terms, aortopathy in males with MFS usually follows a worse course than in females, and male patients usually undergo surgical intervention earlier than females (10–13).

Most of the pathomechanisms identified and subsequent therapies for MFS aortopathy have arisen from the use of mouse models (14–16). The two most popular are the haploinsufficiency/dominant-negative (*Fbn1*^*C1041G/+*^) and the hypomorphic *mgR/mgR* mouse models. These have been widely used to investigate determinant pathomechanisms, such as abnormal cell signaling [mainly transforming growth factor (TGF)-β, extracellular signal-regulated kinase (ERK), and nitric oxide (NO)-associated signaling], hemodynamic and biomechanical forces, redox stress, and extracellular matrix (ECM) remodeling (17–19).

Several new and repurposed pharmacological compounds and gene therapies have been evaluated in these mouse models for their effects on aortopathy progression (both aortic aneurysm and dissection). Unfortunately, very few have entered clinical practice (β-blockers, losartan, and resveratrol), which makes pharmacological therapy complementary, leaving prophylactic aortic surgery as the definitive management, given the high mortality rate after aortic dissection.

Little is known about other factors that accompany the transmission of *FBN1* genetic variants that may influence aortopathy progression. In this sense, it is known that there must be somatic or (epi)genetic modifiers that aggravate or protect patients sharing the same *FBN1* genetic variant. Likewise, it is also well known that the prevalence of the disease is equal, regardless of whether the genetic variant is inherited from the father or the mother (20). Nonetheless, it is unknown whether there is some impact on the cardiovascular manifestations in MFS offspring depending on the parental origin of the *FBN1* pathological variant. If this were the case, this would suggest the presence of some non-genetic parental-associated factor(s) that could also determine, to some extent, the cardiovascular course of the disease in their MFS descendants.

To approach this question initially, we examined some characteristic cardiovascular manifestations of MFS in wild-type (WT) and MFS offspring who inherited the *Fbn1* variant from the father (MFS-P; paternal origin) or the mother (MFS-M; maternal origin). This study aims to provide a first insight into the possible existence of a parental factor(s) that might be transmitted or not, which could impact the offspring, using a simple and well-known animal model of MFS (*Fbn1*^*C1041G/+*^), which presents a uniform genetic background.

## MATERIAL AND METHODS

### 2.1 Ethics

All animal procedures complied with ARRIVE (Animals in Research: Reporting In Vivo Experiments) guidelines and standard ethical regulations (European Union Directive 2010/63/EU). They were approved by the local ethics committee (Animal Experimentation Ethics Committee - University of Barcelona, CEEA-UB) and the Government of Catalonia (347/22).

### 2.2 Animal models

MFS mice carrying a fibrillin-1 mutation (Fbn1C1041G/+; hereafter MFS mice) were obtained from The Jackson Laboratory (B6.129-Fbn1tm1Hcd/J; Strain #012885/Common name: C1039G; Bar Harbor, ME, USA). Both MFS mice and their sex- and age-matched WT littermates were maintained on a C57BL/6J genetic background. All mice were housed, following the institutional guidelines of the University of Barcelona, under controlled conditions (22°C room temperature, 12/12-hour light/dark cycle, 60% humidity) with *ad libitum* access to food and water. Genomic DNA was extracted from a mouse ear punch to determine the genotype of each mouse using PCR. Four experimental groups were generated: WT and MFS control mice (derived from MFS males crossed with WT females) and WT and MFS environment mice (derived from MFS females crossed with WT males). Body weight was measured at multiple time points throughout the study.

### 2.3 Blood pressure measurement

Systolic blood pressure (SBP) was measured in conscious mice using a tail-cuff plethysmography system (Non-Invasive Blood Pressure System, PanLab, Barcelona, Spain). Mice were placed in a restrainer tube, which was cleaned after each use. After an acclimation period, measurements were taken on three separate occasions and averaged for each mouse.

### 2.4 Echocardiography

Two-dimensional transthoracic echocardiography was performed on all animals under 1.5% inhaled isoflurane. Imaging was conducted using a 10–13 MHz phased array linear transducer (IL12i, GE Healthcare, Madrid, Spain) with a Vivid Q system (GE Healthcare, Madrid, Spain). The images were recorded and later analyzed offline using EchoPac software (v.08.1.6, GE Healthcare, Madrid, Spain). Proximal aortic segments were examined in a parasternal long-axis view, and the aortic root diameter was measured from inner edge to inner edge at end-diastole at the level of the sinus of Valsalva. The left ventricular end-diastolic (LVDD) and end-systolic (LVSD) internal diameters were measured in M-mode recordings and were computed by the Teichholz formula into volumes as follows: LVDV=7*LVDD^3^/2.4 (where LVDV is LV end-diastolic volume) and LVSV=7*LVSD^3^/2.4+LVSD (where LVSV is LV end-systolic volume). LV ejection fraction (LVEF) was subsequently calculated as follows: LVEF = ((LVDV-LVSV)/LVDV)*100 and used as a surrogate for LV systolic function. The average of three consecutive cardiac cycles was used for each measurement, with the operator being blinded to the group assignment. Aortic measurements were performed at one, two, four, and six months of age.

### 2.5 Histopathology

Paraffin-embedded tissue arrays of mouse aortae and hearts from all experimental groups were transversely sectioned into 5-µm slices. Aortic sections were stained with Verhoeff-Van Gieson (Merck HT25A), according to the supplier’s instructions, to reveal elastic fibers. Images were captured under visible light using a Leica Leitz DMRB microscope (10X or 20X) equipped with a Leica DC500 camera. All measurements were conducted in a blind manner without knowledge of the genotypes.

Elastic fiber ruptures were quantified by identifying breaks larger than 20 µm (significant discontinuities) in the normal 360° circumferential continuity of each elastic lamina in the aortic media. These breaks were counted in four representative images from three non-consecutive sections of the same ascending aorta.

To analyze vascular smooth muscle cell (VSMC) density and elastic fiber autofluorescence, aortic sections were deparaffinized, rehydrated, stained with DAPI (4’,6-diamidino-2-phenylindole), and mounted on the same glass slide using a fluorescence-compatible mounting medium. Images were acquired using a Widefield Leica AF6000 microscope (40X oil immersion) equipped with a Leica DC500 camera.

### 2.6 Immunofluorescence

Paraffin-embedded cardiac tissue sections were deparaffinized and rehydrated before epitope unmasking. For this, the sections were treated with a retrieval solution (TRIS/EDTA buffer; 5 mL of 1 M Tris-HCl, 1 mL of 0.5 M EDTA, 0.5 mL Tween-20, and distilled H2O to 500 mL) at pH 9 for 30 minutes in a steamer chamber at 95°C. The sections were then incubated with ammonium chloride (NH4Cl, 50 mM, pH 7.4) for 20 min to block free aldehyde groups, followed by permeabilization with 0.3% Triton X-100 for 10 min. After rinsing three times with phosphate-buffered saline (PBS), the sections were incubated with 1% bovine serum albumin (BSA) in PBS for 1 h at room temperature. Next, sections were placed in a humidified chamber overnight at 4°C with the primary antibody (anti-pSMAD2, 1:100; Cell Signaling 3108S or anti-pERK, 1:200; Cell Signaling 9101S; rabbit monoclonal Smad2/3, 1:100, Cell Signaling 8685). The next day, sections were incubated with an anti-rabbit secondary antibody, Alexa 647 (1:1000, Invitrogen). Finally, sections were stained with DAPI and mounted on the same glass slide using a fluorescence-compatible mounting medium. Images were obtained using a Widefield Leica AF6000 equipped with a Leica DC500 camera.

### 2.7 Image Analysis

All images were analyzed using Fiji ImageJ Analysis software (ImageJ 1.54f) (Wayne Rasband and contributors; NIH, USA).

### 2.8 Statistical analysis

Before conducting statistical analyses, the data were tested for normality using the Shapiro-Wilk test. After the relevant test (parametric or not), the data were analyzed to investigate differences due to sex. In those cases where no significant differences were found, the data were grouped only based on genotype. To compare genotypes, Two-way ANOVA with Tukey’s *post-hoc* test or Kruskal-Wallis test with Dunn’s post-test for multiple comparisons was used. To analyze the progression with time, we used a Mixed-effects model (Restricted Maximum Likelihood, REML). Statistical details are shown in the Supplementary Tables. Results are expressed as mean ± SEM (standard error of the mean). A *p*-value less than 0.05 was considered statistically significant. All statistical analyses and graph generation were performed using GraphPad Prism 9 software.

## RESULTS

### Impact on body weight

We first evaluated the weight of WT and MFS offspring of maternal MFS (MFS-M) or paternal MFS (MFS-P) origin (Supplementary Figure 1). There were no statistical differences between WT and MFS offspring (genotype factor) derived from either MFS parent; however, there were statistical differences between males and females (sex factor) at all evaluated ages (one, two, four, and six months) (Supplementary figure 1, panels A, B, C, and D, respectively). Interestingly, one-month-old offspring (both WT and MFS) from MFS-M weighed less than those from MFS-P. However, the evolution with age was just the opposite, where offspring from MFS-M weighed significantly more than offspring from MFS-P. Detailed statistical analysis in Table S1.

### Impact on the aortic root diameter and wall organization

We first evaluated the absolute aortic root (AR) diameter of offspring (WT and MFS; males and females) over time (one, two, four, and six months of age) from MFS-P and MFS-M (Figure 1). No differences were observed between WT and MFS aortae of one-month-old mice when the *Fbn1* mutation came from MFS-P; in contrast, significant differences were evident when the variant came from MFS-M (Figure 1A). Note that the AR diameter of MFS offspring from MFS-M was already significantly larger than that of MFS mice from MFS-P (Figure 1A). At two months of age (Figure 1B), MFS offspring from both MFS-P and MFS-M had a larger AR diameter compared with their respective WT littermates. Interestingly, the AR diameter was significantly larger in both WT and MFS offspring from MFS-M than from MFS-P (Figure 1B), indicating an impact of the MFS-M gestational environment that does not occur in the WT mother (MFS-P). Note that these AR differences between WT and MFS offspring from MFS-P appear at this age, which was not the case when the offspring were one month old (Figure 1E). At the age of four months, the early age-associated difference between MFS offspring from MFS-M and MFS-P was no longer observed (Figure 1C). This is because WT and especially MFS offspring from MFS-P augmented their respective AR diameter faster than WT and MFS offspring from MFS-M (red line in Figure 1E). The statistical difference between WT and MFS offspring genotypes was maintained regardless of the parental origin of the *Fbn1* mutation. No differences in AR diameter were observed when the offspring reached six months of age, regardless of whether they were from MFS-M or MFS-P (Figure 1D), thus consolidating the results previously observed at four months of age. The time course of the AR aneurysm (Figure 1E) shows that the mean baseline AR diameter in MFS offspring from MFS-M (violet line) is higher than in offspring of other genotypes. However, this early difference is progressively mitigated over time (as of two months), and the phenotypic AR difference between WT and MFS genotypes is definitively established, regardless of the maternal gestational environment (WT or MFS). Detailed statistical analysis in Table S2.

**Figure 1.**
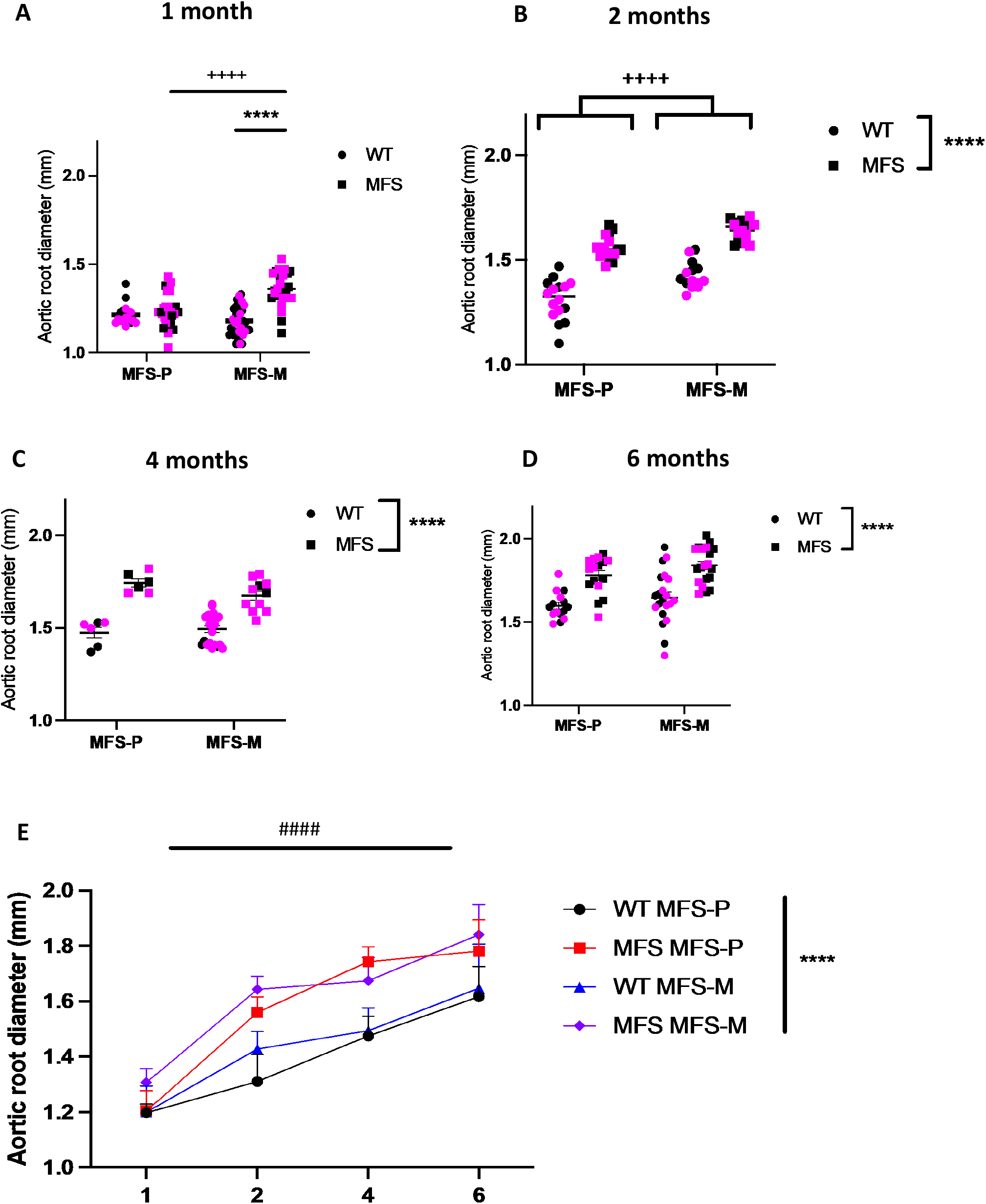
Aortic root diameter over time of offspring with paternal or maternal inheritance of the *Fbn1* genetic variant. Absolute aortic root (AR) diameter evaluated from echocardiographic images in wild-type (WT) and Marfan (MFS) offspring at one, two, four, and six months of age (A-D, respectively) with paternal (MFS-P) or maternal origin (MFS-M) of the *Fbn1* mutation. (E) Overview of AR progression over time. Males are indicated by black points and females by pink points. Results are the mean ± SEM. Statistical analysis: gestational environment (+, MFS-P vs. MFS-M), genotype (*, WT vs. MFS), time (#).

We next studied the histological organization of the aortic wall, evaluating the integrity of elastic fibers and the cellular density of the tunica media (mainly composed of VSMCs) at one and six months of age (Figures 2 and 3, respectively). Concerning elastic fiber integrity, one-month-old MFS offspring from MFS-M had a greater number of large elastic ruptures in the aorta compared with mice from MFS-P, although the difference did not reach statistical significance (Figure 2A). This suggests accelerated MFS aortic wall disarray in MFS offspring from MFS-M. This trend was maintained when evaluating six-month-old mice, where the mean number of elastic ruptures in MFS offspring from MFS-M compared with MFS-P was double (2.8 vs 6, respectively; Figure 2B). Regardless of age, we observed the characteristic genotype differences (WT vs. MFS) in offspring from both MFS-M and MFS-P. Detailed statistical analysis in Table S3.

**Figure 2.**
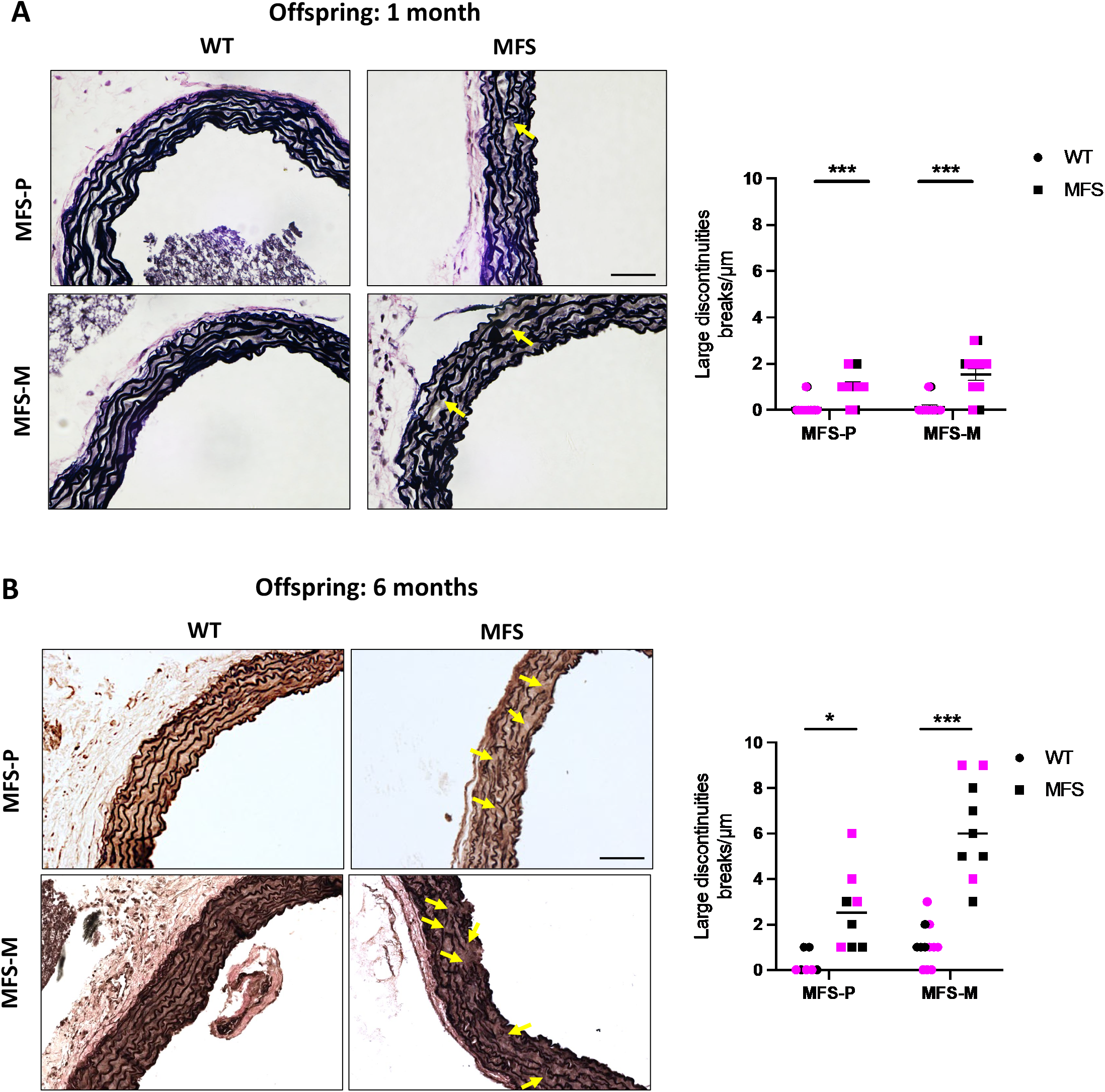
Elastic fiber ruptures in the aortic tunica media of young and adult offspring with paternal or maternal origin of the *Fbn1* genetic variant. Representative images of elastic fibers with Verhoeff-Van Gieson staining from one-(A) and six-month-old aortae (B). Quantitative analysis on the right of each microscopic composition panel. Yellow arrows in A are examples of elastic fiber breaks that were counted. Males are indicated by black points and females by pink points. Results are the mean ± SEM. Statistical analysis: genotype (*, WT vs. MFS).

**Figure 3.**
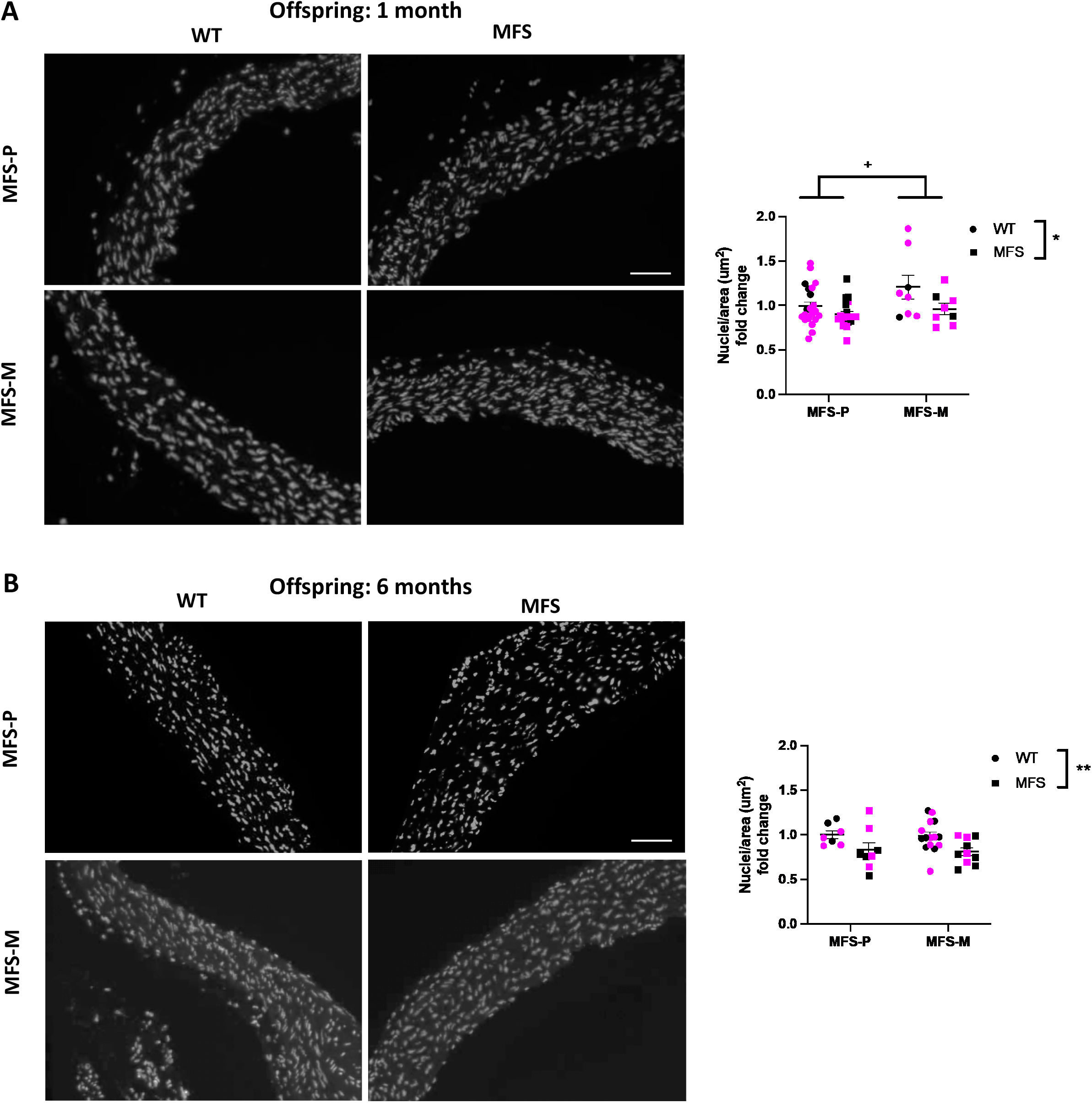
Cell density analysis in the aortic tunica media of young and adult offspring with paternal or maternal origin of the *Fbn1* genetic variant. The cell density in the tunica media was evaluated with DAPI staining of nuclei, both in one-(A) and six-month-old (B) offspring from paternal or maternal MFS (MFS-P and MFS-M, respectively). Males are indicated by black points and females by pink points. Results are the mean ± SEM. Statistical analysis: gestational environment (+, MFS-P vs. MFS-M); genotype (*, WT vs. MFS).

We evaluated the cell density of the tunica media by staining cell nuclei with DAPI. We previously knew that 95% of DAPI-stained cells in the tunica media were also positive for the VSMC marker, α-SMA (Supplementary Figure 2). We observed that one-month-old offspring (WT and MFS) from MFS-M had a greater cell density than mice from MFS-P, particularly in the case of WT offspring (Figure 3A). However, this difference was not observed in six-month-old offspring (Figure 3B). At both ages, MFS offspring exhibited the characteristic MFS-associated cell density reduction. Ddetailed statistical analysis in Table S4.

### Impact on aortic canonical and non-canonical TGF-β signaling

Since TGF-β signaling is a hallmark of the pathogenesis of MFS, we next evaluated canonical and non-canonical pathways by quantifying the number of cells with nuclear immunostaining for the phosphorylated forms of Smad2 (pSmad2) and ERK (pERK) in aortic samples from one- and six-month-old offspring (WT and MFS) from MFS-P or MFS-M. We first analyzed the translocation of pSmad2 to the cell nuclei of the tunica media (Figure 4). In one-month-old offspring (both WT and MFS) from MFS-M, the tunica media presented a greater number of cells with nuclear pSmad2 in comparison with MFS-P offspring (representative images in Figure 4A and quantitative analysis in Figure 4B). This maternal-associated difference disappeared by six months of age (Figures 4A and 4C). At both ages, MFS offspring had a greater number of cells with nuclear pSmad2 compared with WT aortae regardless of the parental origin of the *Fbn1* variant (Figures 4B and 4C). Detailed statistical analysis in Table S5

**Figure 4.**
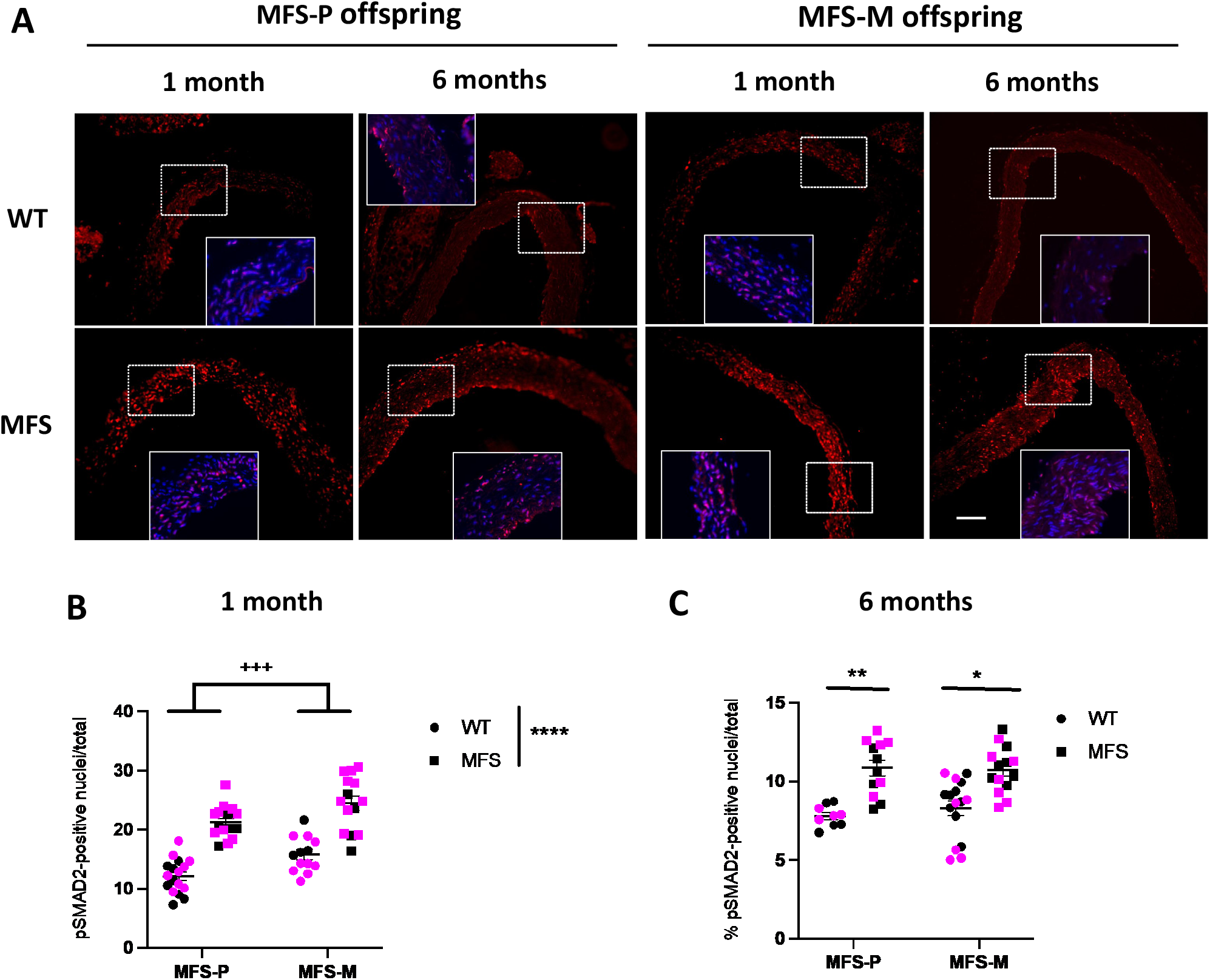
Canonical TGF-β signaling (pSmad2) in young and adult offspring aortae. (A) Immunofluorescence staining for pSmad2 (in red) in the aortae of one- and six-month-old WT and MFS offspring from paternal (MFS-P) or maternal (MFS-M) MFS mice. Insets in A show double staining with DAPI (blue) and pSmad2 (red). (B, C) Quantitative analysis of nuclear pSmad2 for the respective offspring ages. Males are indicated by black points and females by pink points. Results are the mean ± SEM. Statistical analysis: gestational environment (+, MFS-P vs. MFS-M), genotypes (*, WT vs. MFS).

Next, we evaluated the nuclear localization of pERK at both ages (Figure 5). In one-month-old mice, just like for pSmad2, both WT and MFS offspring from MFS-M had a significantly greater number of cells with nuclear pERK than mice from MFS-P (representative images in Figure 5A and quantitative analysis in Figure 5B). As expected, more pERK-stained nuclei were observed in MFS than in WT aortae, regardless of whether the variant came from MFS-P or MFS-M (Figures 5A, 5B, and 5C). Unlike one-month-old offspring, at six months, there were no differences between WT offspring and MFS offspring from MFS-M and MFS-P (Figures 5B and 5C). Detailed statistical analysis in Table S6.

**Figure 5.**
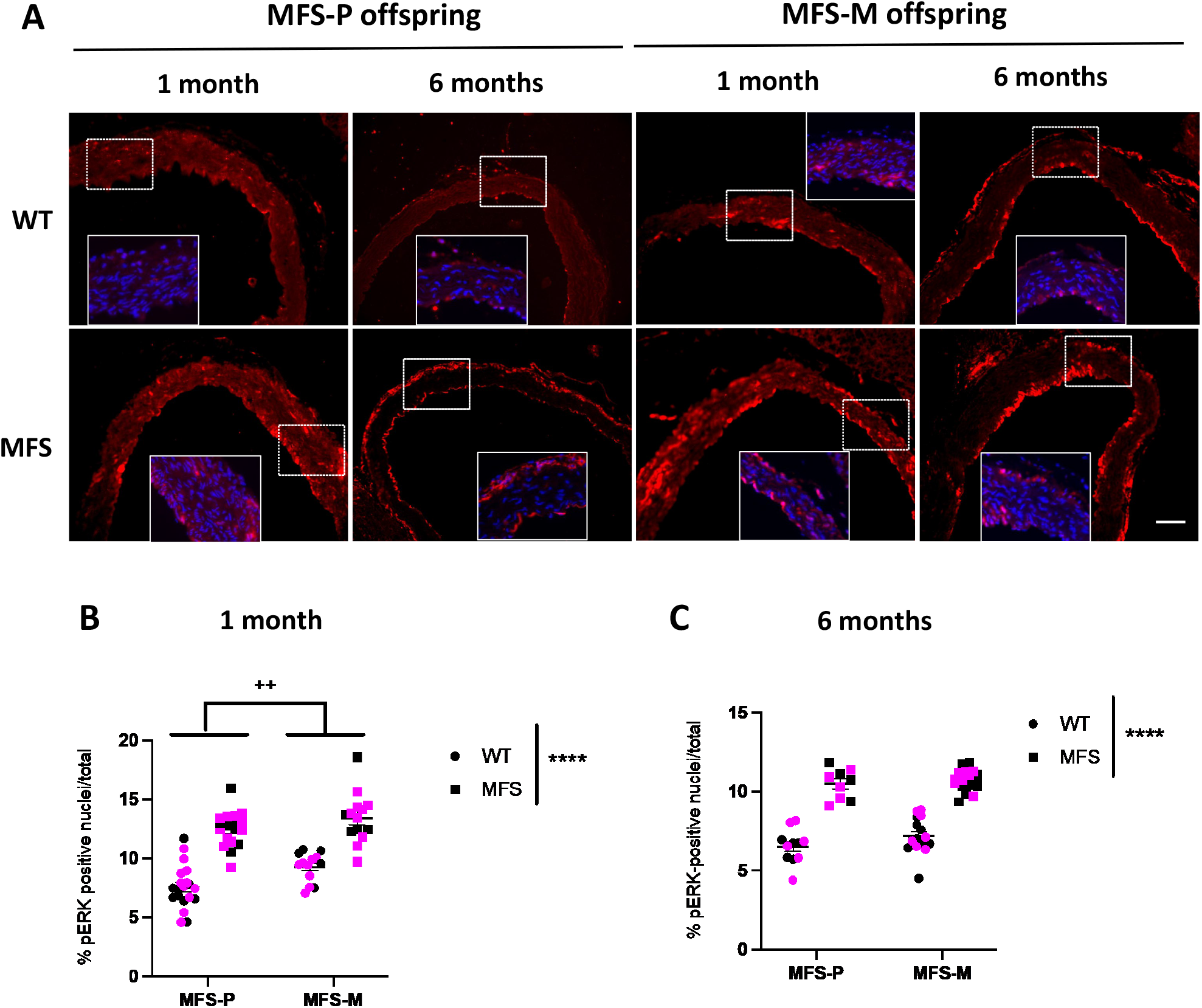
Non-canonical TGF-β signaling (pERK) in young and adult offspring aortae. (A) Immunofluorescence staining for pERK (in red) in the aortae of one- and six-month-old WT and MFS offspring from paternal (MFS-P) or maternal (MFS-M) MFS mice. Insets show double staining with DAPI (blue) and pSmad2 (red). (B, C) Quantitative analysis of nuclear pERK for the respective offspring ages. Males are indicated by black points and females by pink points. Results are the mean ± SEM. Statistical analysis: gestational environment (+, MFS-P vs. MFS-M), genotypes (*, WT vs. MFS).

Interestingly, upon closer examination of the staining patterns of both TGF-β markers, we notice that in one-month-old MFS aortae, both pSmad2 and pERK immunostaining is more pronounced in the tunica media, supporting the idea that early MFS canonical and non-canonical hypersignaling primarily involves VSMCs. However, in six-month-old MFS aortae, both pSmad2 and pERK fluorescence signals were mainly localized in the tunica intima and in the limit between tunicae media and adventitia. Note that, even in six-month-old WT offspring, TGF-β signaling was also primarily localized in the tunica intima, although to a lesser extent than in MFS offspring.

### Impact on the cardiac ejection fraction

We next evaluated the cardiac ejection fraction (EF) (Figure 6), represented by the percentage of blood pumped out by the left ventricle with each heartbeat, indicative of the heart’s capability to pump oxygen-rich blood effectively. In one, two, and four-month-old offspring, there was a significant reduction in EF both in WT and MFS mice from MFS-M compared with those from MFS-P. Regardless of the parental origin of the *Fbn1* mutation, the EF in MFS offspring was consistently significantly lower than in WT animals (*p*=0.0001 at one month; *p*=0.012 at two months, *p*=0.0132 at four months and p=0.0046 at six months). Interestingly, in six-month-old offspring, an interaction was observed between genotype and parental factors. Note that the EF reduction seen in WT and MFS offspring from MFS-M (one, two, and four months of age) remained only in six-month-old MFS offspring (*p*=0.0001, both for MFS vs. WT offspring from MFS-M and MFS offspring from MFS-M vs. MFS-P; Figure 6D). The persistence of EF reduction in six-month-old MFS offspring from MFS-M, but not from MFS-P (p=0.6211), suggests the involvement of a maternal factor that has a lasting impact on adult MFS cardiac function. This effect is absent when the *Fbn1* variant is paternally transmitted and gestation occurs in a WT mother. Detailed statistical analysis in Table S7.

**Figure 6.**
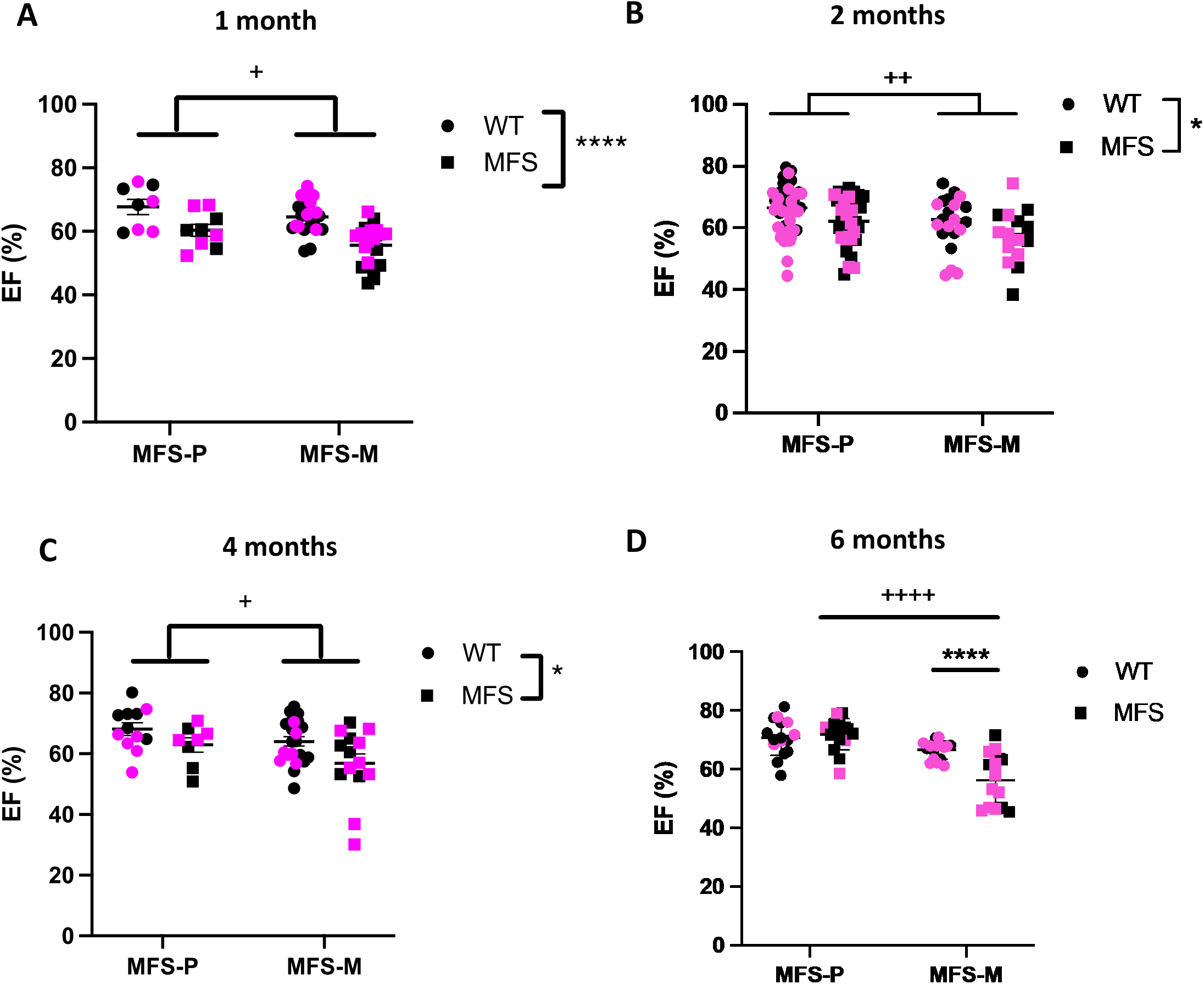
Cardiac ejection fraction over time of offspring with paternal or maternal origin of the *Fbn1* genetic variant. Percentage of ejection fraction (EF) in male and female WT and MFS offspring at different ages (A, one month; B, two months; C, four months; D, six months) from paternal (MFS-P) or maternal (MFS-M) MFS mice. (D) Overview of EF progression over time in the four experimental groups. Results are the mean ± SEM. Males are indicated by black points and females by pink points. Statistical analysis: gestational environment (+, MFS-P vs. MFS-M), genotypes (*, WT vs. MFS).

### Impact on blood pressure

We also evaluated the SBP in offspring at different ages; however, this was only possible in two-, four-, and six-month-old animals, as one-month-old mice were too small for the tail-cuff methodology. Both two-month-old WT and MFS offspring from MFS-M had significantly lower SBP than mice from MFS-P (Figure 7A;). However, this reduction disappeared when mice reached four months of age (Figure 7B). When both WT and MFS offspring from MFS-M reached six months of age, they became hypertensive compared with offspring from MFS-P (Figure 7C). The progressive evolution of the SBP in experimental groups is shown in Figure 7D, where an inverse relationship with age is seen between offspring from MFS-P and MFS-M. Detailed statistical analysis in Table S8.

**Figure 7.**
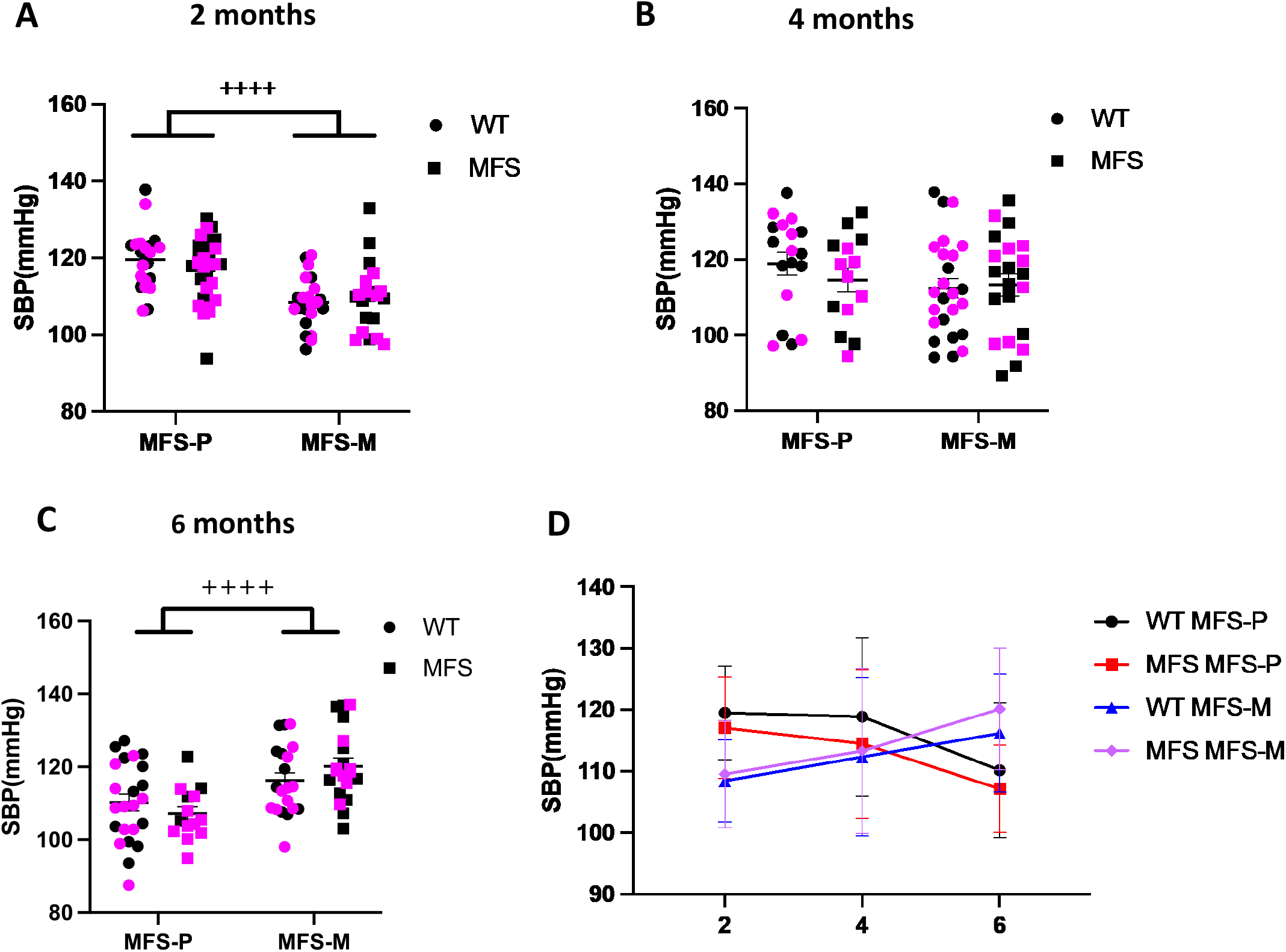
Blood pressure over time of offspring with paternal or maternal origin of the *Fbn1* genetic variant. Systolic blood pressure measurements in WT and MFS offspring from (MFS-P) or maternal (MFS-M) MFS mice at two, four, and six months of age (A-C, respectively). (D) Representation of the temporal course of the SBP in the four experimental groups. Males are indicated by black points and females by pink points. Results are the mean ± SEM. Statistical analysis: gestational environment (+, MFS-P vs. MFS-M).

## DISCUSSION

This study initially aimed to determine whether cardiovascular manifestations in MFS offspring differ depending on the parental origin of the *Fbn1* mutation (MFS-P or MFS-M). However, our findings indicate that, beyond simple parental genetic transmission, the maternal MFS gestational environment exerts a distinct and variable lasting impact on the development of MFS cardiovascular manifestations. We utilized the cysteine mouse model of MFS, *Fbn1*^*C1041G/+*^, as it closely reproduces the typical cardiovascular clinical course of the human disease (18). In general, we observed that MFS-M offspring, both WT and MFS, exhibited a trajectory of adverse cardiovascular and metabolic outcomes, which appeared in early life and persisted in adulthood. Briefly, we found that both WT and MFS offspring from MFS-M were significantly underweight at one month of age, but went on to develop a persistent overweight phenotype from four months onward—an effect not observed in offspring from MFS-P. AR enlargement also presented earlier in MFS offspring from MFS-M and was tightly associated with enhanced TGF-β (Smad2 and ERK) signaling, increased elastic fiber fragmentation, and reduced VSMC density. Notably, early cardiovascular phenotypic differences were also present in WT offspring from MFS-M but were normalized by adulthood. WT offspring from MFS-M exhibited sustained cardiac dysfunction, with the reduced EF persisting into late adulthood, accompanied by a progressive increase in SBP. These findings highlight the existence of an intrauterine gestational environment in MFS mothers that impacts both structural and functional cardiovascular outcomes.

A striking observation was the consistent differences in body weight of both WT and MFS offspring depending on the MFS progenitor. Both WT and MFS offspring from MFS-M, regardless of sex, exhibited a distinct body weight trajectory; while initially underweight at one month of age, they progressively gained weight, resulting in a persistent overweight phenotype by four to six months. Interestingly, the overweight observed in offspring from MFS-M was accompanied by cardiovascular dysfunctions, including an earlier onset of AR dilation in MFS offspring from MFS-M, as well as reduced EF and elevated SBP in both WT and MFS offspring from MFS-M. For instance, nutritional alterations (both restriction and overnutrition) during pregnancy, documented in both animal models and humans, adversely impact cardiovascular development with long-term functional detriment in adulthood (21–27). These effects, present across genotypes and ages, suggest a lasting cardiovascular and metabolic alteration likely driven by an adverse maternal gestational environment, a phenomenon called “fetal programming”. It is well established that a healthy gestational environment is important for the long-term cardiovascular and metabolic health of the offspring (28,29).

### Possible underlying mechanisms

A key mechanistic link in this process is increased maternal oxidative stress, which has been widely implicated in adverse pregnancy outcomes and long-term disease programming (27,30–32). In models of fetal growth restriction, elevated maternal oxidative stress leads to lipid peroxidation and impaired antioxidant defenses in the offspring, altering endothelial function and vascular development (33,34). Exposure to redox stressors during early development may reprogram fetal redox-sensitive pathways, such as those involving NADPH oxidases and mitochondrial metabolism, thus contributing to structural and functional cardiovascular abnormalities. In MFS, increased oxidative stress is a key factor in its pathophysiology, particularly contributing to the development of aortic aneurysms and dissections. Therefore, the maternal environment in our MFS-M model likely induces such oxidative changes during gestation, thereby contributing to fetal programming and the earlier onset of metabolic and cardiac dysfunction in the offspring. While more mechanistic studies are needed, our findings align with a growing body of evidence highlighting maternal oxidative stress as a key mediator of fetal programming, with lasting effects on offspring metabolism, cardiovascular structure, and function.

In the present study, the cardiovascular alterations observed in offspring from MFS-M are likely driven by the early hyperactivation of TGF-β signaling pathways during development. This hypothesis is supported by our finding that MFS offspring from MSF-M exhibited earlier and more pronounced activation of both canonical (pSmad2) and non-canonical (pERK) TGF-β signaling in the aortic wall, detectable in early postnatal stages (one month of age). TGF-β signaling is a central regulator of ECM remodeling, the VSMC phenotype, and inflammation, all processes involved in aortic aneurysm formation in MFS (35). Therefore, the premature activation of these pathways could accelerate elastic fiber fragmentation and VSMC loss, contributing to the early AR dilation observed in MFS-M offspring (36). Moreover, TGF-β signaling is also implicated in cardiovascular remodeling (PMID: 21059352). Sustained activation of pSmad2 and pERK promotes maladaptive cardiac changes, such as reduced contractility and myocardial fibrosis (37), consistent with the persistent decline in EF in MFS-M offspring. Additionally, emerging evidence suggests that TGF-β signaling can influence vascular tone through crosstalk with oxidative stress pathways (see below) and the renin-angiotensin system (38). This could potentially contribute to the progressive increase in SBP and its persistence in adulthood in MFS-M offspring.

An interesting observation revealed changes in TGF-β signaling across postnatal stages in the aortae of MFS offspring. At one month of age, both pSmad2 and pERK signals were mainly seen in the tunica media, while at six months, both signals were predominantly located in the tunica intima in offspring from both MFS-M and MFS-P. This age-associated shift in the main localization of the immunosignal suggests an early overactivation of TGF-β signaling in VSMCs during the initial stages of disease in young MFS offspring, followed by an increased engagement of endothelial cells in later stages of MFS. This is consistent with previous findings showing that TGF-β hypersignaling in MFS aortae alters the VSMC phenotype, shifting from synthetic to contractile (39), and disrupts ECM remodeling, ultimately contributing to the appearance of medial degeneration (35). Histological analyses of human MFS aneurysms similarly demonstrated Smad2/3 accumulation in VSMC and inflammatory cells within the media and subintimal layers (40), with pSmad2 preferentially located in the outer media (41). Our findings at one month align with these observations, indicating that early pathological signaling in MFS primarily involves medial VSMC. The spatial shift in TGF-β signaling in MFS offspring was curiously also seen in WT offspring, although to a lesser extent. At one month of age, the activation of downstream TGF-β signaling was also primarily localized in the tunica intima of WT mice of both parental origins; later, at six months of age, it was more evident in the intima layer, likely reflecting a physiological response to mechanical forces involved in normal vascular growth and ECM remodeling. Notably, this overactivation appeared earlier and was more pronounced in WT offspring from MFS-M compared with MFS-P. This underscores the contribution of the maternal gestational environment to maladaptive cardiovascular aortic remodeling, even in offspring with no genetic predisposition to cardiovascular dysfunction. In contrast, in six-month-old offspring, the dominant localization of pSmad2 and pERK in the intima implies a major shift in signaling activity toward endothelial cells. While many studies have reported that TGF-β signaling is highly active in the media, others have also shown an upregulation of Smad activation in the tunica intima. Endothelial cells, which line the intima layer, express TGF-β receptors and, therefore, can activate Smad signaling in response to pathological stimuli, such as inflammation, altered blood flow, and shear stress. Within this pathological framework, increased TGF-β activity in endothelial cells has been linked to endothelial-mesenchymal transition, vascular inflammation, and fibrotic remodeling (42–44). A plausible mechanism for this spatial shift associated with age and disease progression may involve changes in receptor expression. Recent studies using single-cell RNA sequencing and immunofluorescence in the aortae of patients with MFS reported downregulation of the type II TGF-β receptor in VSMC, despite elevated expression of TGF-β1 ligands (45). If a similar mechanism is at play in the MFS mouse model, the loss of TGF-β receptor expression in medial VSMC may impair downstream responsiveness in this vascular layer. As a result, elevated levels of active TGF-β ligands would increase the signal at neighboring cell types that remain receptive, such as endothelial cells in the tunica intima.

Finally, we also considered the hypothetical contribution of other somatic elements directly involved in the gestational environment of MFS-M, e.g., intrinsic mitochondria-associated dysfunctions and maternal hormonal factors. Consistently, reduced mitochondrial DNA content and oxidative phosphorylation, accompanied by a shift in metabolism toward glycolysis, have been reported in MFS aortic pathology (46). Likewise, it is also probable that our observed cardiac alterations in MFS-M offspring are partially associated with mitochondrial alterations (structure and/or function), with the difference that their impact over time could be greater and/or longer compared with what might occur in the aorta. The hormonal environment during gestation is an additional somatic factor that may contribute to the cardiovascular pathology. Oxytocin—a hormone that initiates uterine contraction and milk letdown throughout lactation—has been reported to enhance the pregnancy-associated risk of aortic dissection in MFS mothers by the hyperactivation of ERK in the aortic wall (47). This study focused solely on gestational mothers; nevertheless, it cannot be discarded that it had an additional impact on the offspring.

## Conclusions

This study demonstrates that maternal inheritance of the MFS *Fbn1* mutation has a significant and lasting impact on the cardiovascular phenotype of the offspring, extending beyond the effects of genetic transmission alone. Notably, some of these alterations were also observed in WT offspring from MFS mothers, reinforcing the notion that the maternal intrauterine environment influences disease trajectory independently of genotype. The cardiovascular alterations and enhanced and spatially dynamic TGF-β signaling observed in WT offspring from MFS-M support a role for gestational programming, potentially driven by maternal oxidative stress and disrupted vascular TGF-β signaling associated with the pathophysiology of maternal MFS. These findings offer new insights into the variability of clinical manifestations in MFS and underscore the importance of considering maternal factors in risk assessment, monitoring, and the development of targeted interventions for children of MFS parents. Finally, conversely to human patients, we did not observed differences between males and females (sex factor) in the here evaluated outcomes.

### Study limitations

We are aware that the study is based on a mouse model of the disease that is widely used to study the pathomechanisms involved in MFS. However, it does not reflect the totality of the significant clinical variability observed in patients; therefore, the conclusions of our study cannot necessarily be translated to human clinical pathology at this moment. This warrants a future study that should be conducted in large patient cohorts.

## Supporting information

Statistic Tables

## Acknowledgments

We are grateful to María Encarnación Palomo for her technical assistance and Helena Kruyer for her accurate editorial assistance.

## Funding

This study was supported by the Spanish Ministry of Science, Innovation and Universities (grant number PID2023-146296OB-I00) and the generous support of patients with Marfan syndrome through private donations and from SIMA (Asociación Española de afectados por el Síndrome de Marfan).

## SUPPLEMENTARY FIGURE LEGENS

Supplementary Figure 1. Body weight of offspring with paternal or maternal inheritance of Marfan syndrome. Both wild type (WT) and Marfan (MFS) offspring from paternal or maternal MFS (MFS-P and MFS-M) were weighed just before echocardiographic measurements and at different ages (A, one month; B, two months; C, four months; D, six months). Males are indicated by black points (left panels) and females by pink points (right panels). Statistical analysis: gestational environment (+, MFS-P vs. MFS-M): genotype (*, WT vs. MFS).

Supplementary Figure 2. Percentage of colocalization in the aortic tunica media (revealed by elastic autofluorescence in green) between nuclei visualized with DAPI (blue points) and the VSMC marker anti-α smooth muscle actin (α-SMA; in red). Dashed white points mark the limit of the media.

## Notes

### Competing Interest Statement

The authors have declared no competing interest.

