## Supplementary figures and images for "Impact of the maternal environment on cardiovascular features of the offspring in a mouse model of Marfan syndrome"

### supplementary figures

#### Slide 1
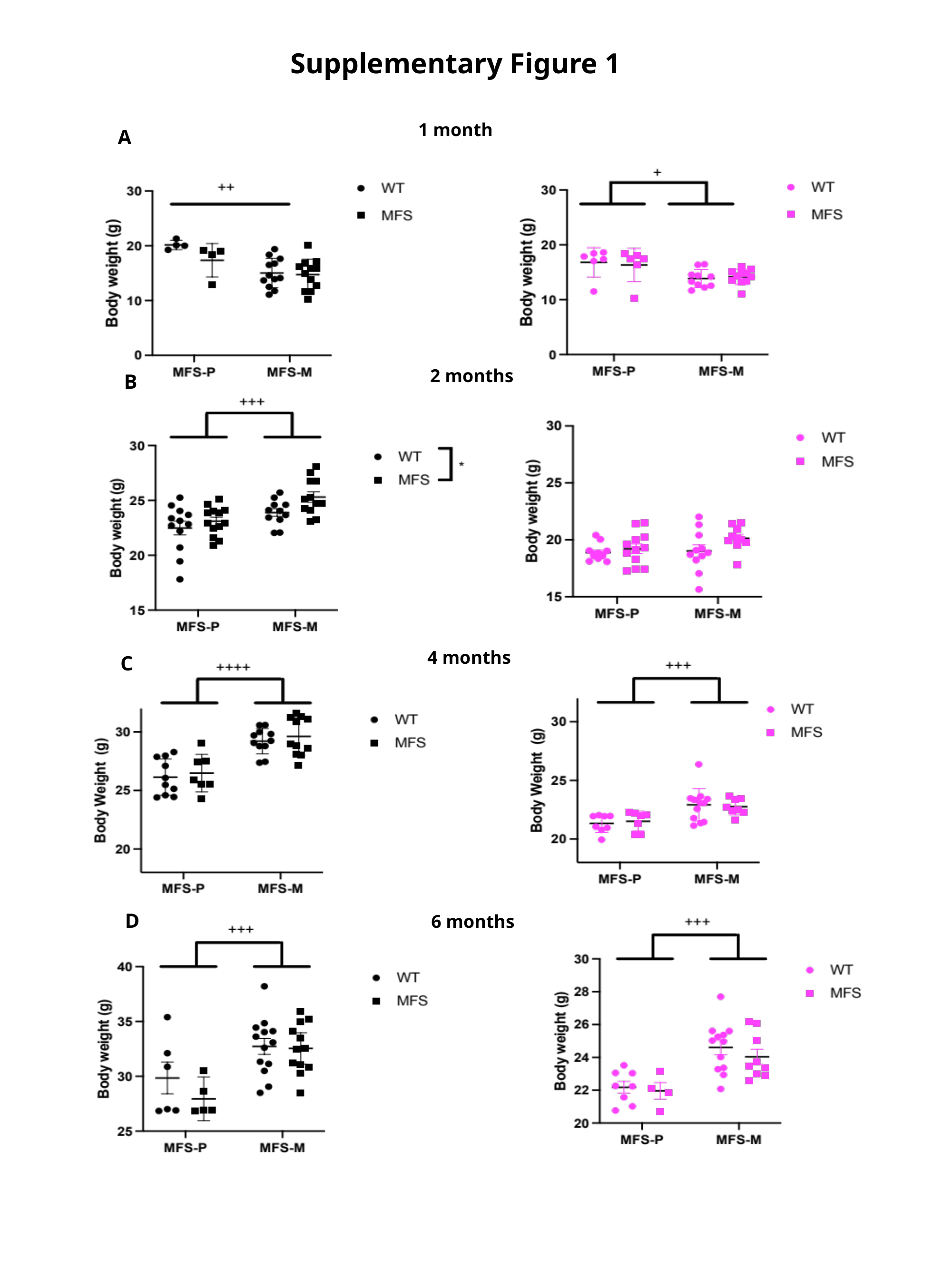

Supplementary Figure 1
1 month
A
2 months
B
4 months
C
D
6 months

#### Slide 2
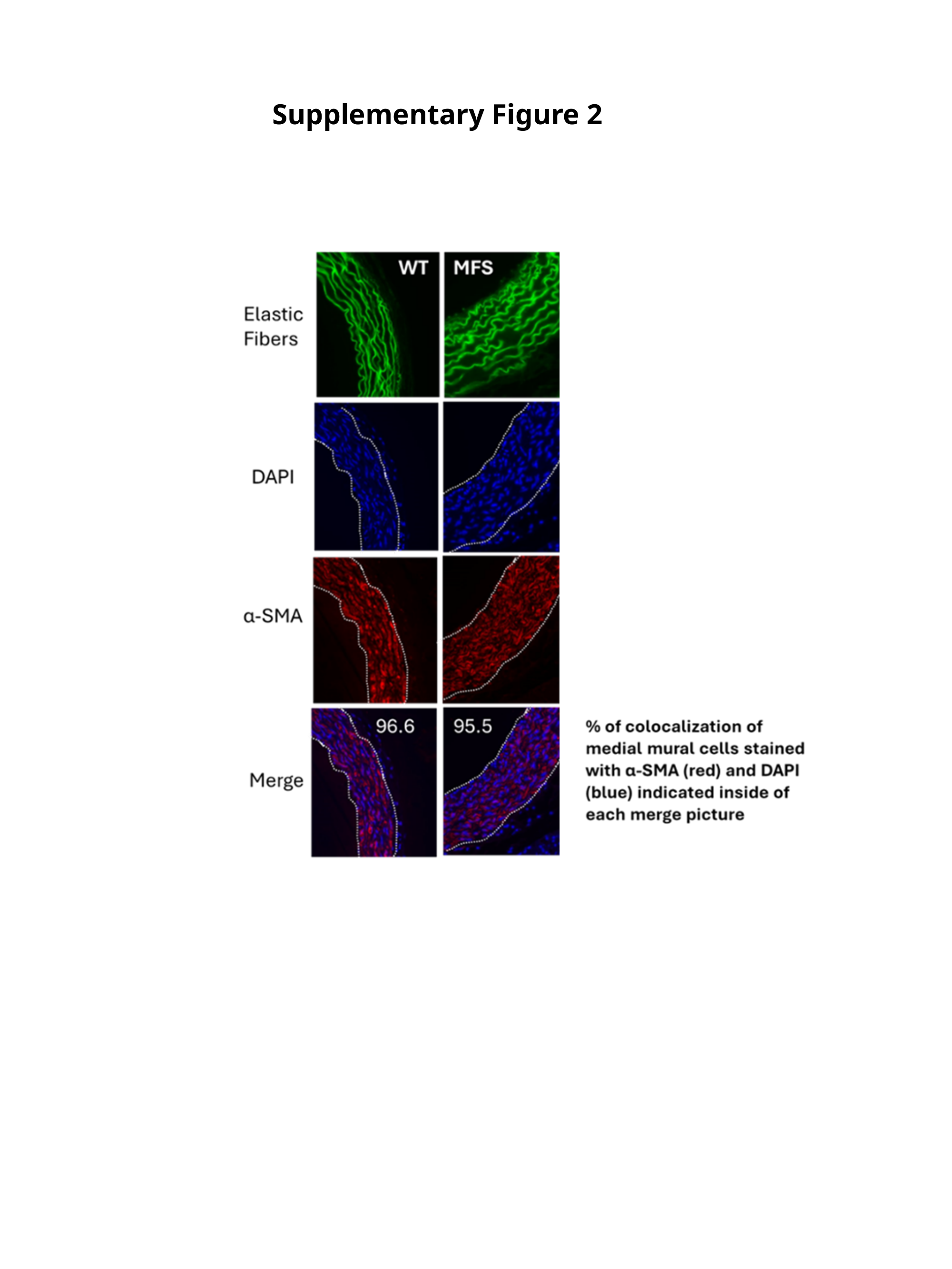

Supplementary Figure 2
